# The *Tomato brown rugose fruit virus* movement protein overcomes *Tm-2*^*2*^ resistance while attenuating viral transport

**DOI:** 10.1101/2020.12.13.420935

**Authors:** Hagit Hak, Ziv Spiegelman

## Abstract

*Tomato brown rugose fruit virus* (ToBRFV) is a new virus of the *Tobamovirus* genus, causing substantial damage to tomato crops in the Middle East. Reports of recent ToBRFV outbreaks from around the world indicate an emerging global epidemic. ToBRFV overcomes all tobamovirus resistances in tomato, including the durable *Tm-2*^*2*^ resistance gene. Here, we show that the ToBRFV movement protein (MP^ToBRFV^) is the cause for overcoming *Tm-2*^*2*^ resistance. Transient expression of MP^ToBRFV^ failed to activate the *Tm-2*^*2*^ resistance response. Replacement of the original MP sequences of *Tomato mosaic virus* (ToMV) with MP^ToBRFV^ enabled this recombinant virus to overcome *Tm-2*^*2*^ resistance. Hybrid protein analysis revealed that the resistance-breaking elements are located between MP^ToBRFV^ amino acids 1 and 216, and not the C terminus as previously assumed. Interestingly, replacement of *Tobacco mosaic virus* (TMV) and ToMV MPs with MP^ToBRFV^ caused an attenuation of systemic infection of both viruses. Cell-to-cell movement analysis revealed that MP^ToBRFV^ moves less effectively compared to the TMV MP (MP^TMV^). These findings suggest that overcoming *Tm-2*^*2*^ is associated with attenuated MP function. This viral fitness cost may explain the high durability of *Tm-2*^*2*^ resistance, which had remained unbroken for over 60 years.

## 1. Introduction

The *Tobamovirus* genus is one of the most devastating groups of plant viruses, which includes the *Tobacco mosaic virus* (TMV), *Tomato mosaic virus* (ToMV) and *Cucumber green mottle mosaic virus* (Broadbent, 1976; Rybicki, 2015; Dombrovsky *et al.*, 2017). Tobamoviruses form rod-shaped virions, which encompass a single-stranded, sense RNA genome of about 6.4 kb. This genome encodes four main open reading frames (ORFs): Two subunits of a 126-kDa protein and its read-through 183 kDa variant forming the viral replication complex, a 30-kDa movement protein (MP) and a 17.5 kDa coat protein (CP) (Goelet *et al.*, 1982; Ishibashi and Ishikawa, 2016). Cell-to-cell movement of the virus depends on the viral-encoded MP, which binds to multiple cellular components to transport the viral RNA through plasmodesmata, membrane-lined channels interconnecting adjacent cells (Lucas, 2006; Ueki and Citovsky, 2011; Pitzalis and Heinlein, 2018; Reagan and Burch-Smith, 2020).

A recent outbreak of a new tobamovirus named *Tomato brown rugose fruit virus* (ToBRFV) has substantially damaged the tomato industry in Israel and Jordan, leading to extensive reductions in both yield and fruit quality (Salem *et al.*, 2016; Luria *et al.*, 2017). Recent reports of ToBRFV outbreaks around the world indicate an emerging global epidemic (Cambrón-Crisantos *et al.*, 2018; Yan *et al.*, 2019; Menzel *et al.*, 2019; Ling *et al.*, 2019; Skelton *et al.*, 2019; Fidan *et al.*, 2019; Panno *et al.*, 2019). ToBRFV overcomes all known tobamovirus resistances in tomato, including the durable *Tm-2*^*2*^ resistance (*R*) gene, and therefore poses a substantial threat for the global tomato industry (Luria *et al.*, 2017).

Dominant resistance (*R*) genes are currently the most powerful means to control plant pathogens (de Ronde *et al.*, 2014). Most of them encode nucleotide-binding leucine-rich repeat (NLR) class of intracellular immune receptors (DeYoung and Innes, 2006). NLRs recognize specific pathogen-encoded effectors either directly or indirectly (Padmanabhan and Dinesh-Kumar, 2014) and trigger an immune signaling cascade that confines the pathogen to the infection site, typically through one of two processes: induction of programmed cell death (PCD) in a process termed hypersensitive response (HR) or extreme resistance, in which resistance is achieved apparently without cell death. Most plant NLRs contain a coiled-coil (CC) or Toll-interleukin-1-receptor (TIR) homology domain at their N-terminus, a centrally located nucleotide binding (NB) domain, and a leucine rich repeat (LRR) domain at the C-terminus (Burdett *et al.*, 2019).

Resistance against tobamoviruses in tomato was achieved by the introgression of the *R* genes *Tm-1*, *Tm-2* and its allele *Tm-2*^*2*^ from wild tomato species (Meshi *et al.*, 1989; Ishibashi *et al.*, 2007; Lanfermeijer*et al.*, 2003). In recent decades, resistance-breaking strains have emerged for *Tm-1* and *Tm-2*, leaving *Tm-2*^*2*^ as the key gene for control of tobamovirus in tomato for the last 60 years (Meshi *et al.*, 1989; Strasser and Pfitzner, 2007). *Tm-2*^*2*^ encodes a CC-NLR, which associates with its effector, tobamovirus MP, on the plasma membrane to activate an immune response (Weber *et al.*, 1993; Chen *et al.*, 2017). This association confers plant resistance through the induction of either HR or extreme resistance (Hall, 1980), an outcome which depends on the specific level of *Tm-2*^*2*^ expression (Zhang *et al.*, 2013). Recently it was shown that, upon activation by the MP, Tm-2^2^ proteins self-associate on the plasma membrane to form an oligomer that relays the immune signal in a process that depends on both its CC and NB domains (Wang *et al.*, 2020). Previous studies concluded that the C-terminal part of MP is essential for recognition by Tm-2^2^ (Weber *et al.*, 1993; Weber and Pfitzner, 1998); two amino-acid substitutions at the ToMV MP (MP^ToMV^) C terminus of the resistance-breaking ToMV isolate ToMV-2^2^, S238R and K244E, were found to overcome *Tm-2*^*2*^ resistance (Weber *et al.*, 1993). Moreover, a premature stop codon in MP^ToMV^, causing truncation of the MP C-terminal 30 amino acids, enabled ToMV to overcome *Tm-2*^*2*^ resistance (Weber and Pfitzner, 1998).

ToBRFV is currently the only known tobamovirus that overcomes *Tm-2*^*2*^ resistance (Luria *et al.*, 2017). Previous sequence analysis showed that ToBRFV contains 21 potential resistance-breaking mutations. Of those, 12 mutations are located in the ToBRFV MP (MP^ToBRFV^) and likely contribute to overcoming *Tm-2*^*2*^ (Maayan *et al.*, 2018). These include mutations that potentially change protein features such as secondary structures, phosphorylation sites and disulfide bonds. However, so far, there was no experimental evidence for the role of MP^ToBRFV^ in overcoming *Tm-2*^*2*^ resistance.

Here, we show that MP^ToBRFV^ overcomes *Tm-2*^*2*^ resistance, independent of its C-terminal part. In addition, we show that overcoming *Tm-2*^*2*^ is associated with reduced systemic and cell-to-cell movement, suggesting that the resistance-breaking elements have negative effects on MP^ToBRFV^ function. These findings may explain the high durability of *Tm-2*^*2*^, which has only been broken recently by ToBRFV.

## 2. Results

### 2.1 MP^ToBRFV^ does not trigger *Tm-2*^*2*^-mediated cell death

Activation of NLR receptors by their effectors can be monitored by the appearance of HR-mediated cell death when the NLR is co-expressed with its effector (Whitham *et al.* 1996; Moffett *et al.*, 2002; Wu *et al.*, 2015). Multiple studies showed that co-expression of *MP*^*TMV*^ and *Tm-2*^*2*^ can trigger HR in *N. benthamiana* (Zhang *et al.*, 2013; Chen *et al.*, 2017; Wang *et al.*, 2020). To test if Tm-2^2^ recognizes MP^ToBRFV^, the coding sequences of MP^ToBRFV^ and MP^TMV^ were cloned under the control of the *35S* promoter (*p35S:MP*^*ToBRFV*^ and *p35S:MP*^*TMV*^, respectively). Clones were then transiently expressed in leaflets of tomato cv. Ikram (a *Tm-2*^*2*^*/tm-2* heterozygote) using agroinfiltration (Figure 1 a-c). When *p35S:MP*^*TMV*^ was expressed, cell death was observed in infiltrated leaflets as a result of immune response activation (Figure 1a). On the other hand, leaflets infiltrated with *p35S:MP*^*ToBRFV*^ showed only marginal necrosis (Figure 1b), similar to that of the empty vector control treatment (Figure 1c). To confirm that these findings are a direct result of *Tm-2*^*2*^ activity, a similar experiment was performed with *N. benthamiana*. Here again, transient co-expression of *p35S:MP*^*TMV*^ with *p35S:Tm-2*^*2*^ caused marked lesions and electrolyte leakage consistent with the establishment of HR (Figure 1d and i), while co-expression of *p35S:MP*^*ToBRFV*^ with *p35S:Tm-2*^*2*^ resulted in minimal cell death (Figure 1e) and reduced electrolyte leakage (Figure 1i). Individual expression of *Tm-2*^*2*^, *MP*^*ToBRFV*^ and *MP*^*TMV*^ resulted in no cell death or electrolyte leakage (Figure 1f-i). The marginal immune activation by MP^ToBRFV^ indicated that this protein was able to evade *Tm-2*^*2*^ resistance.

**Figure 1.**
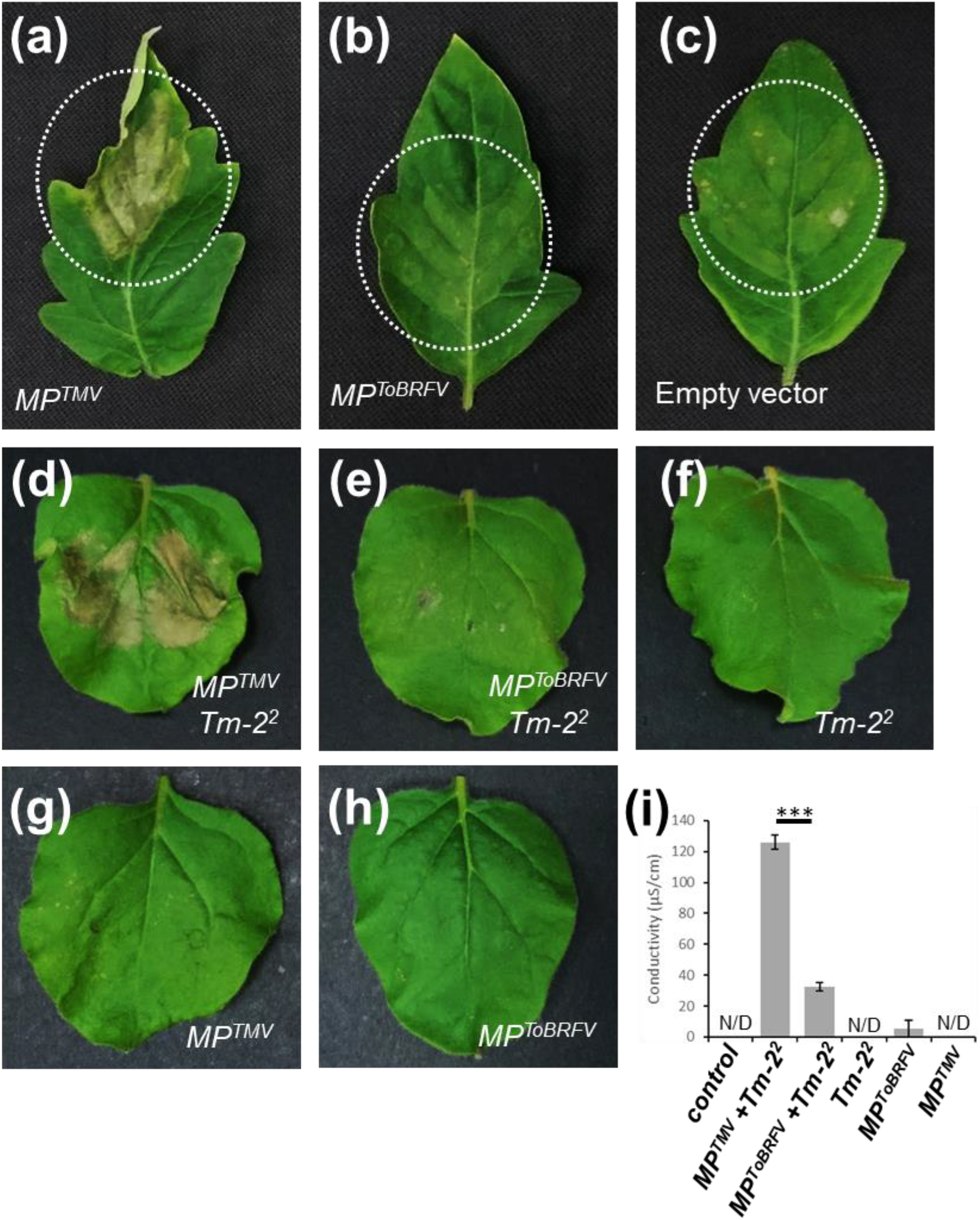
*Tm-2*^*2*^-mediated cell death in tomato and *N. benthamiana* in response to MP^TMV^ and MP^ToBRFV^. (a-c) Transient expression of *MP*^*TMV*^ (a), *MP*^*ToBRFV*^ (b) and and empty vector in tomato cv. Ikram containing the Tm-22 resistance gene. (d-h) Transient expression of *MP*^*TMV*^ with *Tm-2*^*2*^ (d), MP^ToBRFV^ with *Tm-2*^*2*^ (e), *Tm-2*^*2*^ (f), *MP*^*TMV*^ (g) and MP^ToBRFV^ (h) in *N. benthamiana* leaves. (i) Electrolyte leakage assay of *N. benthamiana* leaves expressing the various constructs (***P<0.001, student’s t-test, *n* ≥ 4). N/D = not detected.

### 2.2 A recombinant clone of ToMV harboring MP^ToBRFV^ overcomes *Tm-2*^*2*^ resistance in tomato

To test the ability of MP^ToBRFV^ to overcome *Tm-2*^*2*^ resistance in tomato, a hybrid clone was generated based on an existing infectious clone of ToMV (Hamamoto *et al.*, 1993). The original MP sequence of ToMV was replaced with MP^ToBRFV^ (ToMV^MP-ToBRFV^) (Figure 2a). This clone was then used to infect tomato plants homozygous for the ToMV-sensitive *tm-2* allele or the *Tm-2*^*2*^ allele (Figure 2b). Viral RNA was quantified in infected and systemic leaves (Figure 2c and d, respectively) using RT-qPCR. As expected, inoculation of *tm-2* plants with ToMV resulted in severe viral symptoms (Figure 2b) and high level of viral accumulation in both infected and systemic leaves (Figure 2c-e), while *Tm-2*^*2*^ plants were completely immune to ToMV (Figure 2b-e). In marked contrast, ToMV^MP-ToBRFV^ induced symptoms in both *tm-2* and *Tm-2*^*2*^ plants (Figure 2b and e) and viral accumulation was detected in plants harboring both alleles (Figure 2c-e). It is important to note that in both *tm-2* and *Tm-2*^*2*^ ToMV^MP-ToBRFV^ symptoms were milder than those of ToMV-infected *tm-2* plants (Figure 2b), suggesting that MP^ToBRFV^ reduced ToMV pathogenicity. Together, these results established that MP^ToBRFV^ is sufficient to overcome *Tm-2*^*2*^ resistance.

**Figure 2.**
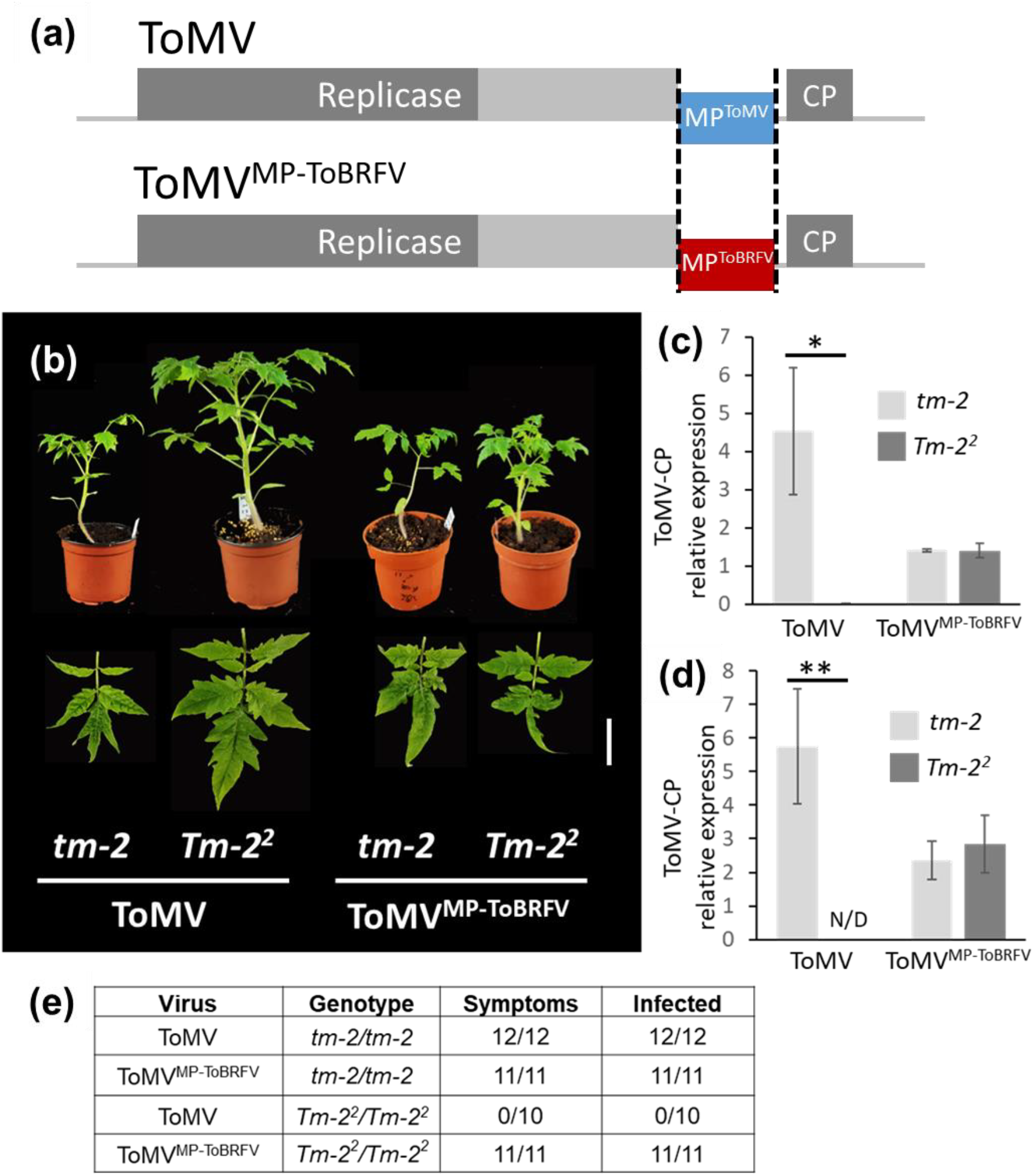
Replacement of MP^ToMV^ with MP^ToBRFV^ is sufficient to overcome *Tm-2*^*2*^ resistance in tomato. (a) Schematic illustration of ToMV and ToMV^MP-ToBRFV^ viruses. Segmented lines indicate the replacement of MP^ToMV^ (blue) with the MP^ToBRFV^ (red) sequence. (b) Tomato plants (cv. Moneymaker) (upper panel) and leaves (lower panel) homozygous to the *tm-2* or *Tm-2*^*2*^ allele infected with ToMV and ToMV^MP-ToBRFV^. (c-d) RT-qPCR analysis for ToMV coat protein (CP) transcripts in locally infected (c) and systemic (d) leaves of *tm-2* and Tm-2^2^ tomato plants infected with ToMV and ToMV^MP-ToBRFV^. (e) Number of plants showing systemic symptoms of ToMV infection in and results of virus detection by RT-PCR assay (*P<0.05, **P<0.01, student’s t-test, *n* ≥ 4). N/D = not detected. Scale bar = 2 cm.

### 2.3 The MP^ToBRFV^ C terminus is dispensable for overcoming Tm-2^2^

To characterize the potential resistance-breaking elements in MP^ToBRFV^, a protein sequence alignment was performed on MP^ToBRFV^, MP^TMV^, MP^ToMV^ and the MP of the *Tm-2*^*2*^ resistance-breaking ToMV isolate ToMV-2^2^ (MP^ToMV-2^2^^) (Figure 3a). At the whole protein level, MP^ToBRFV^ was 81% identical to MP^ToMV^ and 78% identical MP^TMV^. However, at the C-terminal 50 amino acids (216-266) (Figure 3a, broken line), the sequence was more variable (41% and 42% identity with MP^ToMV^ and MP^TMV^, respectively). Importantly, this region includes the C-terminal 30 amino acids and MP^ToMV^ amino acids S238 and K244, previously shown to be essential for recognition by *Tm-2*^*2*^ (Figure 3a, yellow) (Weber and Pfitzner, 1993; Weber *et al.*, 1998). This suggested that the MP^ToBRFV^ C terminus is the resistance-breaking factor.

**Figure 3.**
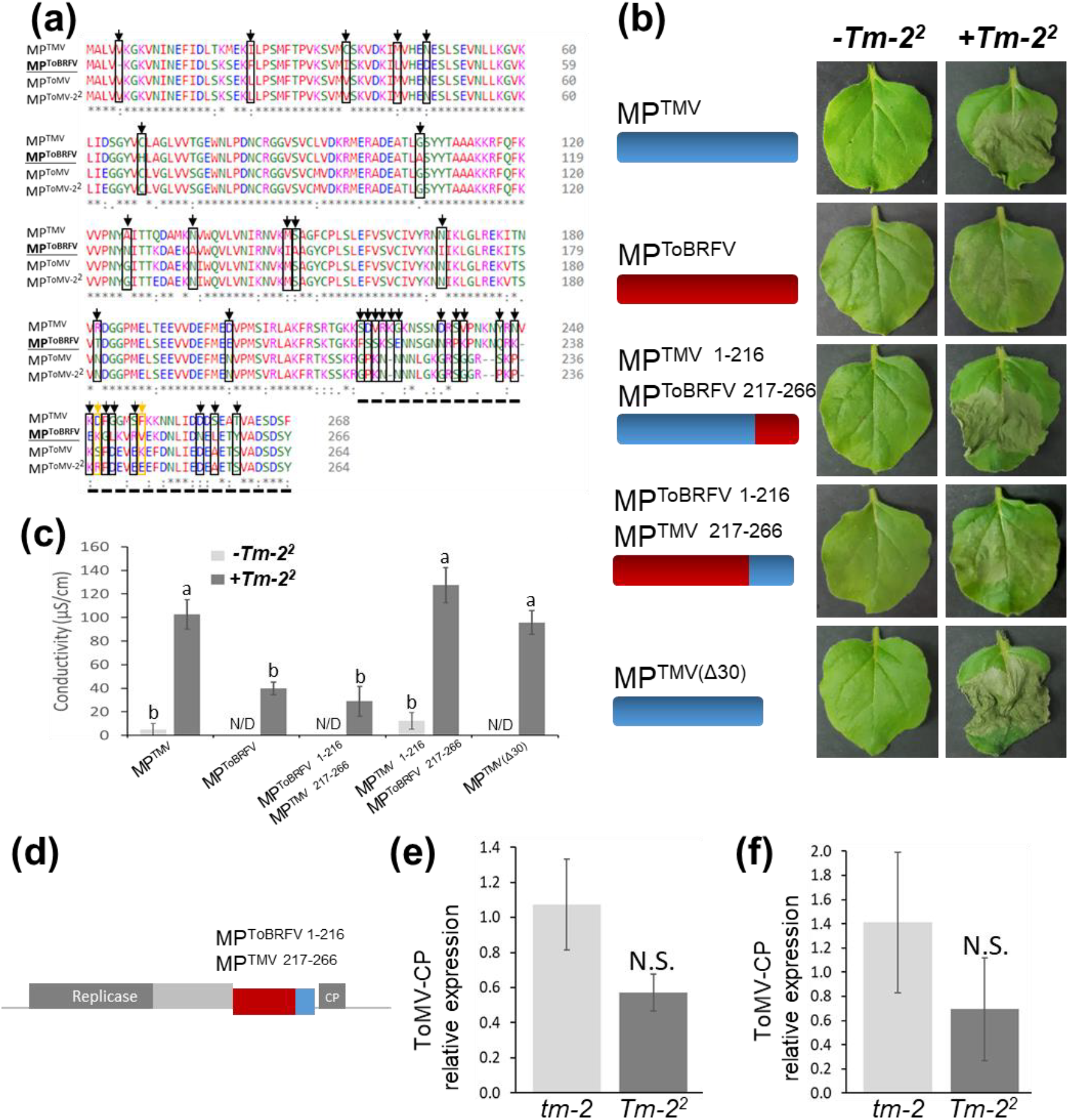
Overcoming *Tm-2*^*2*^ resistance is not due to the MP^ToBRFV^ C terminus. (a) Amino acid sequence alignment of MP^ToBRFV^ with MP^TMV^, MP^ToMV^ and the resistance-breaking MP^ToMV-22^. A total of 33 suspected resistance-breaking mutations are highlighted in black. The MP^ToMV^ amino acids Ser238 and Lys244, previously found to be required for Tm-2^2^ recognition, are highlighted in yellow. The segmented black line underlies the variable 50 C-terminal amino acids. (b) Transient expression of MP^TMV^, MP^ToBRFV^, the hybrids MP^ToBRFV 1-216^/MP^TMV 217-266^ and MP^TMV 1-216^/MP^ToBRFV 217-266^ and the truncated MP^TMV(Δ30)^ in *N. benthamiana* leaves. Clones were expressed alone (−*Tm-2*^*2*^) or co-expressed with *Tm-2*^*2*^ (+*Tm-2*^*2*^). (c) Electrolyte leakage assay of *N. benthamiana* leaves expressing the various constructs. (d) Schematic illustration of the ToMV^MP-ToBRFV 1-216^ virus expressing a hybrid movement protein with MP^ToBRFV^ amino acids 1-216 fused to MP^TMV^ amino acids 217-266 (e-f) RT-qPCR analysis for ToMV coat protein (CP) transcripts in infected (e) and systemic (f) leaves of tm-2 and *Tm-2*^*2*^ tomato plants infected with ToMV^MP-ToBRFV 1-216^. Different letters indicate significant in Tukey’s HSD test. (*n* ≥ 4). N.S. = not significant.

To test if the MP^ToBRFV^ C terminus overcomes *Tm-2*^*2*^, two hybrid MP clones were generated (Figure 3b): the first containing MP^TMV^ amino acids 1-216 fused the MP^ToBRFV^ C-terminal 50 amino acids 217-266 (MP^TMV 1-216^/MP^ToBRFV 217-266^) and the second containing MP^ToBRFV^ amino acids 1-216 fused to MP^TMV^ amino acids 217-266 (MP^ToBRFV 1-216^/MP^TMV 217-266^) (Figure 3b). Co-expression of *p35S:Tm-2*^*2*^ with MP^TMV 1-216^/MP^ToBRFV 217-266^ induced strong HR, indicating that the MP^ToBRFV^ C terminus was insufficient to evade *Tm-2*^*2*^ (Figure 3b and c). In addition, the MP^ToBRFV 1-216^/MP^TMV 217-266^ fusion did not trigger cell death when co-expressed with *Tm-2*^*2*^ (Figure 3b and c). To verify that the MP^TMV^ C-terminal region is not required for *Tm-2*^*2*^ recognition, a clone with a truncated version of MP^TMV^ containing a stop codon in position 238 was generated (MP^TMV Δ30^). Here again, a marked HR was observed when MP^TMV Δ30^ was co-expressed with *Tm-2*^*2*^, suggesting that the MP^TMV^ C-terminus is dispensable for *Tm-2*^*2*^ recognition (Figure 3b and c).

To confirm these results in tomato, a hybrid ToMV clone was generated with MP^ToBRFV^ amino acids 1-216 fused to MP^ToMV^ amino acids 217-266 (Figure 3d). Viral transcripts were detected in infected and systemic leaves of both *tm-2* and *Tm-2*^*2*^ plants 3 weeks after inoculation (Figure 3e and f), establishing that this virus was able to overcome *Tm-2*^*2*^ resistance, regardless of the MP^ToBRFV^ C terminus.

### 2.4 MP^ToBRFV^ attenuates systemic viral infection of TMV-GFP and ToMV

Overcoming dominant resistance is often associated with viral fitness costs (García‐Arenal and Fraile, 2013). Since MP^ToBRFV^, which overcomes *Tm-2*^*2*^ resistance, also caused reduced symptoms in *tm-2* tomato plants (Figure 3b), we hypothesized that the function of MP^ToBRFV^ is reduced as compared to other MPs. To test this, a TMV-GFP clone (Lindbow, 2007) harboring MP^ToBRFV^ instead of its original MP (TMV-GFP^MP-ToBRFV^) was generated. Local and systemic infection were determined 7 days after inoculation (dpi) of *N. benthamiana* plants using GFP *in vivo* imaging analysis (IVIS) (Figure 4). While TMV-GFP showed strong fluorescence in both locally infected and systemic leaves (Figure 4a, d-f), the resistance-breaking TMV-GFP^MP-ToBRFV^ hybrid showed weaker fluorescent signals (Figure 4b, d-f). To confirm that the reduced infection was due to the attenuated function of MP^ToBRFV^, a rescue assay was done by co-agroinfiltrating TMV-GFP^MP-ToBRFV^ with *p35S:MP*^*TMV*^. Interestingly, MP^TMV^ partially rescued the effect of MP^ToBRFV^ on systemic spread of the virus (Figure 4c and d), leading to a 2-fold increase in local infection and a 4-fold increase in systemic accumulation of the virus (Figure 4e and f, respectively). These results suggested that MP^ToBRFV^ is less effective than MP^TMV^ in promoting viral spread, likely due to its reduced function in cell-to-cell transport of the virus.

**Figure 4.**
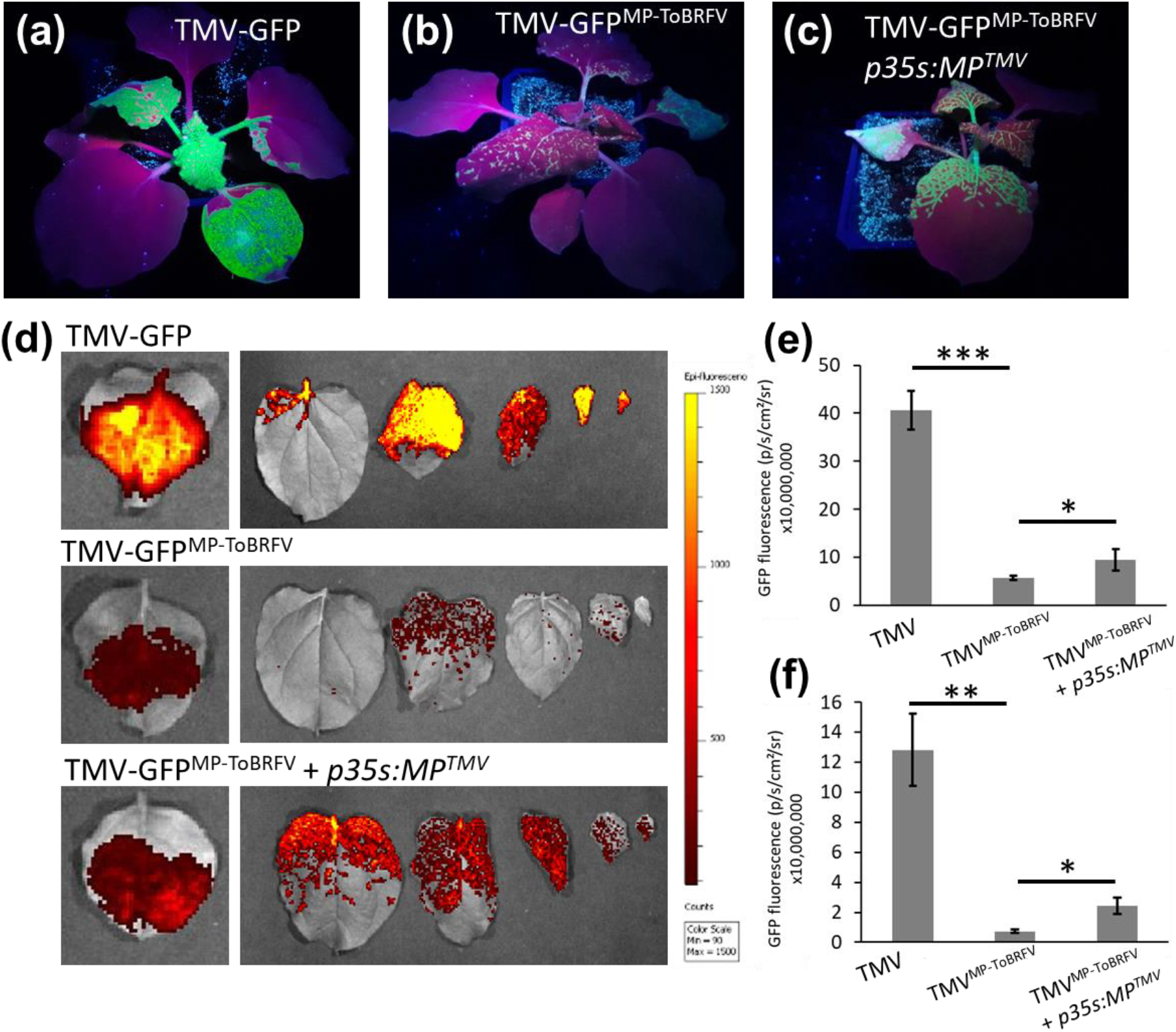
Replacement of MP^TMV^ with MP^ToBRFV^ causes a reduction in TMV-GFP systemic infection in *N. benthamiana*. (a-c) UV images of *N. benthamiana* plants infected with TMV-GFP (a), TMV-GFPMP-ToBRFV (b) and TMV-GFP^MP-ToBRFV^ co-expressed with *p35S:MP*^*TMV*^ (c). (d) IVIS images of infected (left panel) and systemic leaves 1-4 from the apex (right panel) of infected plants. (d-e) Quantitative analysis of GFP fluorescence in locally infected (e) and systemic (f) leaves of plants expressing TMV-GFP , TMV-GFP^MP-ToBRFV^ and TMV-GFP^MP-ToBRFV^ co-expressed with *p35S:MP*^*TMV*^. (*P<0.05, **P<0.01, student’s t-test, *n* ≥ 5).

We further monitored the effect of MP^ToBRFV^ on systemic spread in *tm-2* tomato plants by western blot analysis of CP accumulation (Figure 5). Time course analysis of virus accumulation was performed in systemic tomato leaves 4, 8 and 12 dpi of plants infected with ToMV, ToMV^MP-ToBRFV^ and ToBRFV (Figure 5a). ToMV was first detected in systemic leaves at 4 dpi, while the viruses harboring MP^ToBRFV^, ToMV^MP-ToBRFV^ and ToBRFV, were barely detected. At 8 dpi, ToBRFV and ToMV^MP-ToBRFV^ could be detected in systemic leaves, however, their signal was significantly weaker than ToMV (Figure 5a and b). At 12 dpi there was no significant differences in accumulation of all three viruses. These results suggest that, while MP^ToBRFV^ confers the ability to overcome *Tm-2*^*2*^, it also reduces the rate of systemic infection compared to MP^TMV^ and MP^ToMV^.

**Figure 5.**
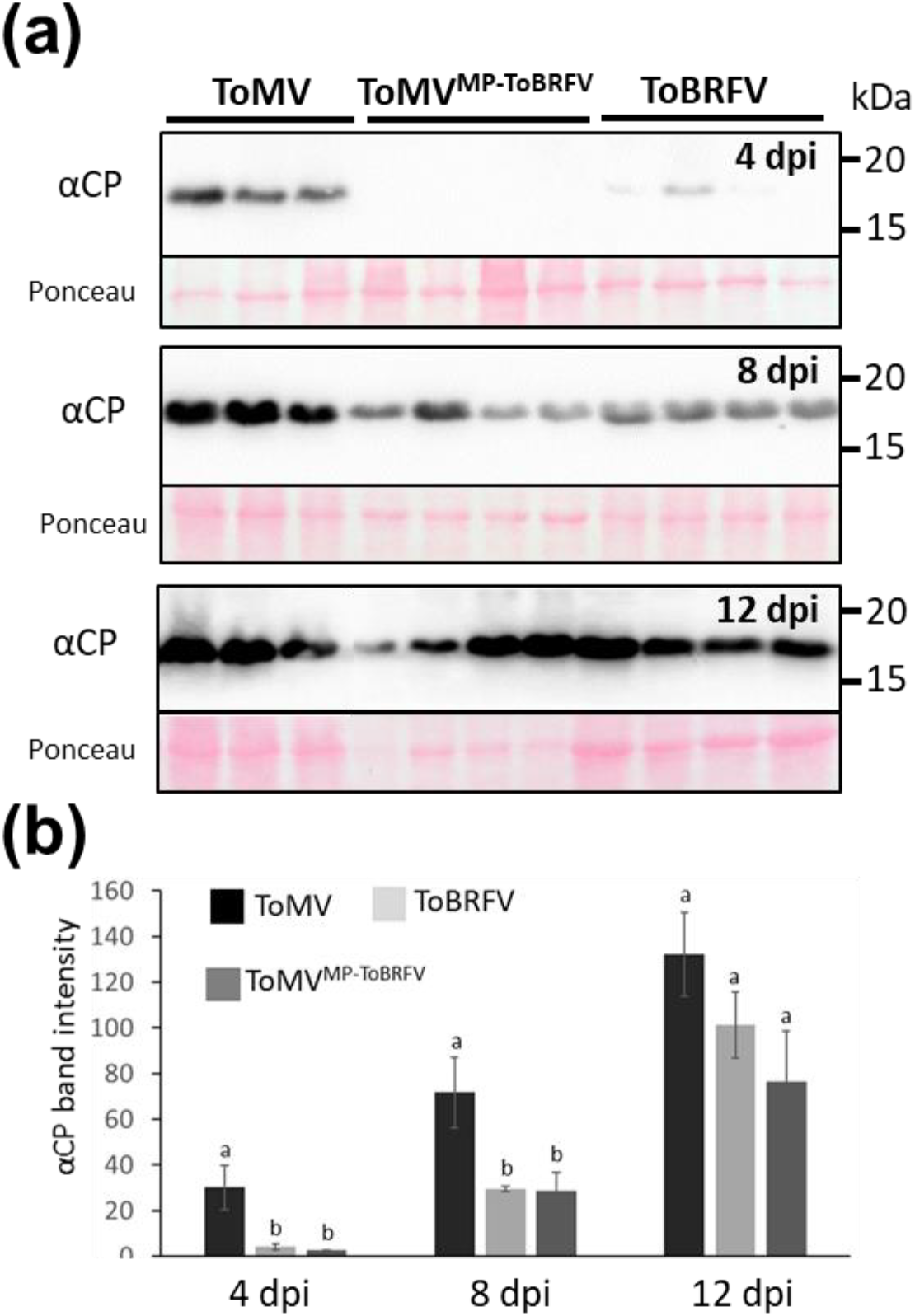
Delayed systemic infection by viruses expressing MP^ToBRFV^ in tomato plants. (a) Western blot analysis of ToMV, ToMV^MP-ToBRFV^ and ToBRFV coat protein in systemic leaves 4, 8 and 12 days post inoculation (dpi). (b) Quantitative analysis of ToMV, ToMV^MP-ToBRFV^ and ToBRFV coat proteins in systemic leaves. Note that while ToMV coat proteins are detected in systemic leaves at 4 dpi, ToBRFV and ToMV^MP-ToBRFV^ coat proteins accumulate later, at 8 dpi. Different letters indicate significantly different values (P <0.05, Tukey’s HSD test, n ≥ 4).

### 2.5 Reduced cell-to-cell transport of MP^ToBRFV^ compared to MP^TMV^

Since the role of a MP is to enable intercellular transport of the viral RNA, we speculated that the reduction of systemic infection caused by MP^ToBRFV^ is a result of a decrease in cell-to-cell movement. To monitor cell-to-cell movement, the ORFs of both *MP*^*TMV*^ and *MP*^*ToBRFV*^ were fused to *YFP* and expressed under the *35S* promoter (Figure 6a and d) in *N. benthamiana* leaves. As an immobile control, ER-targeted mCherry (ER-mCherry) was expressed from the same binary plasmid, to mark the original cell of expression (Figure 6b and e). Low dilution (OD = 0.0001) of agrobacterium was used to express the constructs in individual *N. benthamiana* epidermal cells and z-stack confocal images were taken for each cell. The result was cell clusters, in which the central cell expressed the MP and the proximal cells contained the trafficked MP (Figure 6c and f). The number of adjacent cells containing the trafficked MP^TMV^-YFP and MP^ToBRFV^-YFP was used to quantify the level of intercellular movement. Interestingly, while MP^TMV^-YFP moved to an average of 9.28±0.38 cells (*n* = 64), MP^ToBRFV^-YFP moved to an average of 6.8±0.38 cells (*n* = 51) (Figure 6g). In addition, movement to ≥10 cells was evident in 48.4% of MP^TMV^-YFP cell clusters, but only in 13.2% of MP^ToBRFV^-YFP cell clusters. Consistently, movement to ≤4 cells was detected in 25.5% of MP^ToBRFV^-YFP but only 6.25% of MP^TMV^-YFP (Figure 6h). These results show that MP^ToBRFV^ intercellular movement is reduced as compared to MP^TMV^.

**Figure 6.**
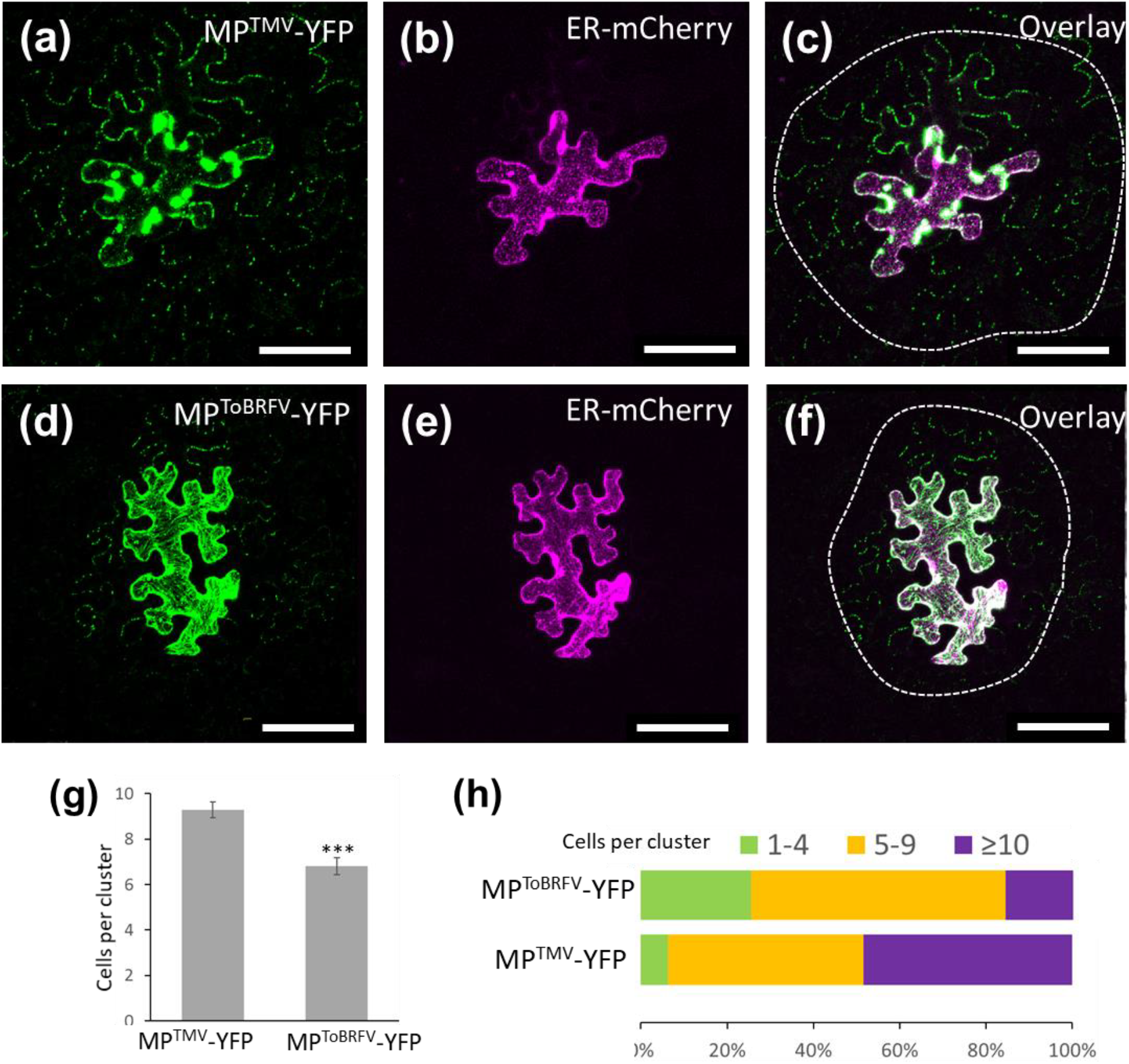
Reduced cell-to-cell movement of *MP*^*ToBRFV*^ compared to *MP*^*TMV*^. (a-c) Coexpression of MP^TMV^-YFP (a) and the non-mobile ER-mCherry (b) in *N. benthamiana* leaf epidermal cells. (c) Overlay of MP^TMV^-YFP and ER-mCherry. (d-f) Coexpression of MP^ToBRFV^-YFP (d) and the non-mobile ER-mCherry (e) in N. benthamiana leaf epidermal cells. (f) Overlay of MP^ToBRFV^-YFP and ER-mCherry. Broken white lines circle the cluster of cells to which the MPs moved. (g) Quantitative analysis of the number of cells per MP^TMV^-YFP and MP^ToBRFV^-YFP clusters. (h) Fractionation of MP^TMV^-YFP and MP^ToBRFV^-YFP clusters: 1-4 cells (green), 5-9 cells (yellow) and ≥10 (purple). (***P < 0.001, Student’s t-test n ≥ 51). Scale bar = 100 μm.

## 3. Discussion

ToBRFV is a significant threat to tomato production worldwide. This highly infectious virus can be easily transmitted by seed coats, working hands, tools, soil and beneficial pollinators (Smith and Dombrovsky, 2019; Levitzky *et al.*, 2019; Klap *et al.*, 2020a). In addition, interaction of ToBRFV with other viruses may increase their disease severity (Luria *et al.*, 2018; Klap *et al.*, 2020b). The main cause for the global ToBRFV outbreak is that this novel virus overcame the long-standing *Tm-2*^*2*^ resistance. Since *Tm-2*^*2*^-based resistance remained unbroken for decades, the question of which factors caused this resistance to break is of agricultural and scientific importance. Previous studies established that the viral MP is recognized by Tm-2^2^ to trigger resistance against the virus. Therefore, in the current study, we investigated the role of MP^ToBRFV^ in overcoming *Tm-2*^*2*^. Our findings establish that *Tm-2*^*2*^ is not activated by MP^ToBRFV^ (Figure 1), and that MP^ToBRFV^ is sufficient to overcome *Tm-2*^*2*^ (Figure 2). These findings are consistent with previous findings on the role of the viral MP as the target of Tm-2^2^ resistance and establish MP^ToBRFV^ as the *Tm-2*^*2*^ resistance-breaking factor (Meshi *et al.*, 1989; Weber *et al.*, 1993).

While it has been shown that the C-terminal part of MP is essential for recognition by Tm-2^2^ (Weber *et al.*, 1993; Weber and Pfitzner, 1998), later studies suggested that the C-terminal part of the MP is dispensable for recognition by Tm-2^2^. Expression of a truncated version of MP^ToMV^, consisting of only the N-terminal 188 amino acids, was sufficient to trigger the *Tm-2*^*2*^ immune response, while expression of the C-terminal 76 amino acids alone did not activate *Tm-2*^*2*^ resistance (Weber *et al.*, 2004). In addition, a TMV MP (MP^TMV^) variant with a C-terminal 81 amino acid deletion still activated *Tm-2*^*2*^ resistance in *N. benthamiana* (Chen *et al.*, 2017). Here, we showed that the MP^ToBRFV^ C-terminal part does not determine *Tm-2*^*2*^ recognition or avoidance. Instead, it seemed that elements within MP^ToBRFV^ amino acids 1-216 overcome this resistance. Computational analysis of MP^ToBRFV^ revealed 12 potential resistance-breaking amino acids, which are not conserved with MP^TMV^ and MP^ToMV^ and can alter the protein charge, structure or phosphorylation, many of which are located between amino acids 1 and 216 (Maayan *et al.*, 2018). Our study provides a platform for elucidating the precise MP^ToBRFV^ amino acids are required to overcome *Tm-2*^*2*^.

Overcoming host resistance is often associated with reductions in viral fitness (García‐Arenal and Fraile, 2013). There are numerous examples in which resistance-breaking mutations cause penalties in viral infectivity or competition. For example, resistance-breaking mutations in the *Potato virus Y* Nla protein, which overcome the *R* gene *Ry*, also caused loss of its protease activity (Mestre *et al.*, 2003). Mutations in *Turnip mosaic virus* which overcome TuRB01 and TuRB04 resistances in *Brassica napus* impaired the virus’s ability to compete with a non-resistance-breaking viral strain on a susceptible host (Jenner *et al.*, 2002). ToMV replicase mutations that overcome *Tm-1* resistance also cause an attenuation of viral replication (Ishibashi *et al.*, 2012). Recently, it was shown that resistance-breaking mutations in the *Pepper mild mottle virus* coat protein variously affect viral particle stability (Bera *et al.*, 2019). Here, we show that overcoming *Tm-2*^*2*^ resistance by MP^ToBRFV^ is associated with delayed viral spread and cell-to-cell movement (Figures 4 to 6).

Previous reports suggested that overcoming *Tm-2*^*2*^ might require mutations that impair viral movement or survival, and that existing *Tm-2*^*2*^ resistance-breaking isolates display reduced transmissibility (Hall, 1980). Indeed, MP^ToBRFV^ amino acids 1-216 include elements essential for MP function (Maayan et al., 2018), including plasmodesmal localization signal (Yuan *et al.*, 2016), a region essential for increasing plasmodesmal size exclusion limit (Weigmann *et al.*, 1994), endoplasmic reticulum interaction domain (Peiró *et al.*, 2014) and RNA binding domains (Citovsky *et al.*, 1992). In addition, the MP C-terminal 50 amino acids, which are dispensable for overcoming *Tm-2*^*2*^ (Figure 3), are also non-essential for viral movement (Berna *et al.*, 1991). The finding that MP^ToBRFV^ activity is reduced compared to MP^TMV/ToMV^ supports the hypothesis that *Tm-2*^*2*^ targets MP sequences essential for its function.

In light of these findings, one may wonder how ToBRFV was able to break this resistance, whereas TMV and ToMV failed to do so. To address this question, we propose the hypothetical model outlined in Figure 7. According to this model, resistance-breaking mutations are required for MP^ToMV^ or MP^TMV^ to overcome Tm-2^2^ (Figure 7a). These resistance-breaking mutations would likely impair MP function leading to loss of viral movement, and eventually loss of infectivity or pathogenicity (Figure 7b). Reconstitution of viral movement would require additional compensatory mutations for MP to retain functionality (Figure 7c). The requirement of multiple mutations to overcome *Tm-2*^*2*^ combined with high levels of fitness penalties would be highly unlikely or rare and uncompetitive. ToBRFV, on the other hand, likely evolved on an unknown host (Maayan *et al.*, 2018), and invaded tomato due to a host shifting event (Figure 7d). Evolution on another host allowed MP^ToBRFV^ to diverge from MP^TMV^ and MP^ToMV^, in a manner that will allow it to evade *Tm-2*^*2*^, yet sustain sufficient levels of movement (Figure 7d).

**Figure 7.**
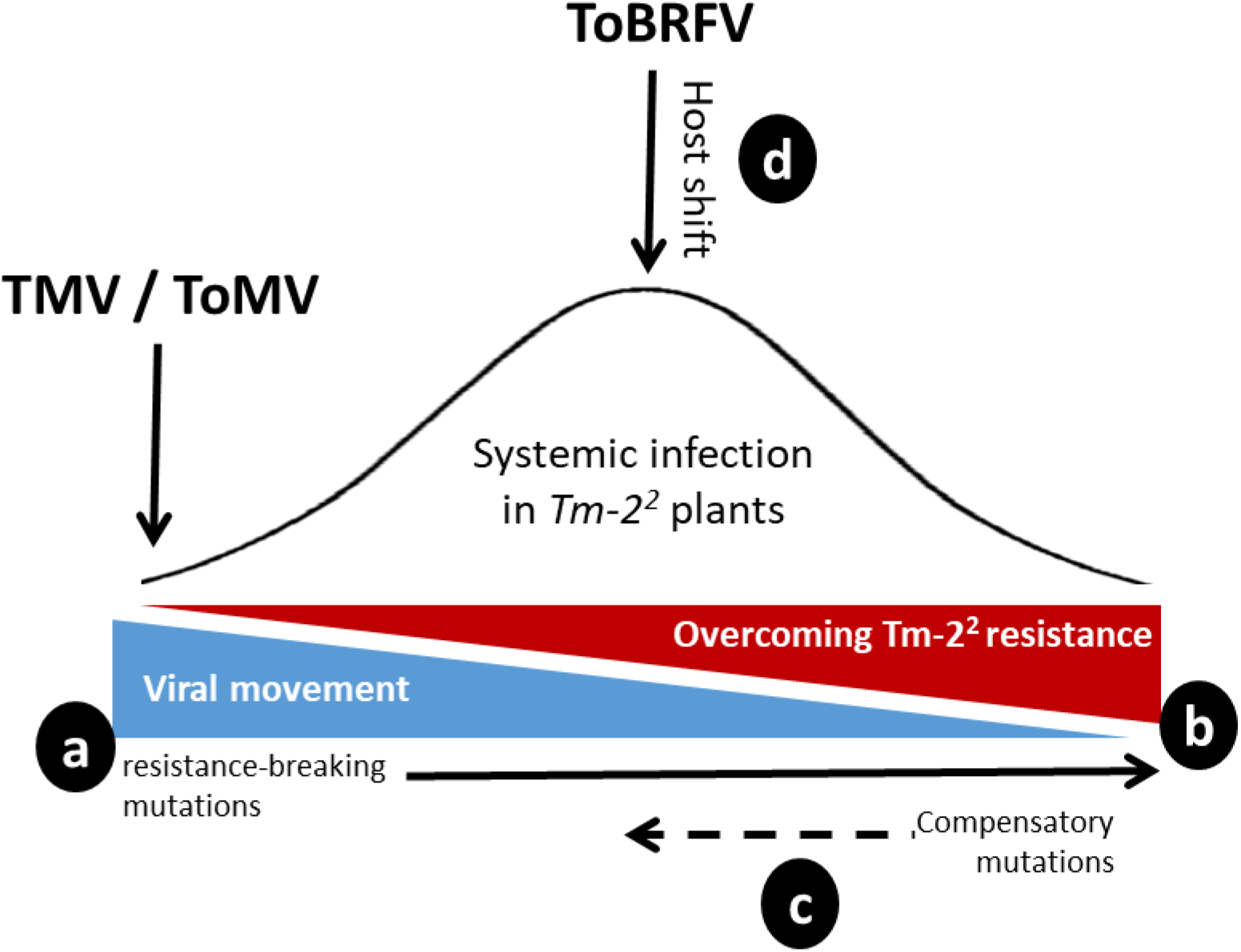
Hypothetical model for overcoming *Tm-2*^*2*^ resistance. (a) To overcome *Tm-2*^*2*^ resistance, MP^TMV^ or MP^ToMV^ will develop resistance-breaking mutations, which will likely impair their movement capacity (b). (c) To restore MP function, multiple compensatory mutations would be required. Such processes, which require multiple rounds of mutation and selection, will be unlikely to occur. (d) ToBRFV, whose emergence in tomato is likely a result of a host shift, accumulated sufficient mutations, which enable it to function while not being recognized by Tm-2^2^.

Collectively, our results establish MP^ToBRFV^ as the *Tm-2*^*2*^ resistance-breaking factor. Based on our systemic and cell-to-cell movement assays, we propose that the mutations that prevented activation of Tm-2^2^ may have also affected MP function. This may explain the high durability of *Tm-2*^*2*^ against TMV and ToMV. We conclude that the MP^ToBRFV^ protein sequence balances between *Tm-2*^*2*^ evasion and viral movement.

## 4. Experimental procedures

### 4.1 Plant materials

Tomato *(Solanum lycopersicum* L.) seeds (cv. Moneymaker) homozygous for the *tm-2* allele (LA2706) or the *Tm-2*^*2*^ allele (LA3310) were obtained from the Tomato Genome Research Center (TGRC), University of California, Davis. Tomato plants cv. Ikram were obtained from Syngenta Co. *Nicotiana benthamiana* and tomato plants were grown in soil in a light- and temperature-controlled chamber at 25°C with a 16h light/8h dark regime. Five-week-old *N. benthamiana* plants were used for agroinfiltration of TMV-GFP and three-week-old tomato plants were used for mechanical inoculation of ToMV.

### 4.2 Plasmid Construction

The various MP^TMV^/MP^ToBRFV^ hybrids were generated using fusion PCR. The following binary plasmids were assembled by Golden Gate cloning using the MoClo Tool Kit for plants (Addgene kit #1000000044) (Weber *et al.*, 2011). *p35S:Tm-2*^*2*^, *p35S:MP*^*TMV*^ and *p35S:MP*^*ToBRFV*^ were generated by cloning the *Tm-2*^*2*^ (AF536201), MP^TMV^ and MP^ToBRFV^ ORFs into pICH486988, a binary Level 2 Golden Gate vector that contains an expression cassette under the CaMV *35S* promoter. To allow Golden Gate assembly, internal BsaI sites were removed by one-step cloning. Plasmids containing the double expression cassettes *p35S:MP*^*TMV*^*-YFP; ER-mCherry* and *p35S:MP*^*ToBRFV*^*-YFP; ER-mCherry* were assembled using the Level 2 vector pAGM4673. The ER-mCherry sequence were amplified from the *ER‐rk CD3‐959* plasmid (Nelson *et al.*, 2007), containing the mCherry ORF with the *Arabidopsis thaliana wall‐associated kinase 2* signal peptide at the N‐terminus of the ER His‐Asp‐Glu‐Leu retention signal at its C‐terminus.

pTLW3 is a cDNA vector harboring the full ToMV sequence under the T7 promoter (Hamamoto *et al.*, 1993). ToMV^MP-ToBRFV^ was assembled by fusion PCR and the MP^ToBRFV^ ORF was inserted between the KpnI and XmaI sites of the vector. The binary vector pJL24 (Lindbow, 2007) contains the full TMV genome with the GFP gene upstream to the MP ORF, and is named here TMV-GFP. The MP^ToBRFV^ ORF was amplified from RNA extracted from ToBRFV-infected plants (accession no. KX619418.1) and was cloned between the AgeI and PacI sites of the pJL24 vector using fusion PCR to create TMV-GFP^MP-ToBRFV^.

### 4.3 Transient expression and agro-infiltration

*Agrobacterium tumefaciens* strain EHA105 harboring a binary vector were grown overnight at 28°C. Cell cultures were resuspended in MES buffer (10mM MgCl2, 10mM MES, 150μM acetosyringone, pH 5.6) to an optical density of OD_600_ = 0.5 (or OD600 = 0.0001 for cell-to-cell movement experiments) and infiltrated using a needleless syringe into the abaxial side of the 4th or 5th leaf of *N. benthamiana.*

### 4.4 Electrolyte leakage assay

*p35S:Tm-2*^*2*^ and *p35S:MP* constructs were agroinfiltrated at OD_600_=0.25 individually or at a 1:1 ratio as indicated to *N. benthamiana* leaves. Twenty-seven hours later, four leaf discs (1 cm diameter) were cut out of each leaf and washed for 10-30 minutes in distilled water. The leaf discs were then carefully transferred into a tube containing 4 ml of distilled water and placed in a growth chamber. Electric conductivity was measured after 6 hours using a benchtop conductivity meter.

### 4.5 TMV-GFP infection and quantification

*A. tumefaciens* harboring TMV-GFP or TMV-GFP^MP-ToBRFV^ vectors was infiltrated into the fourth leaf of 4-week-old *N. benthamiana* plants. GFP fluorescence was monitored 6 days post infiltration. Images were acquired and analyzed using In Vivo Imaging System (IVIS Lumina LT, PerkinElmer) equipped with a XFOV-24 lens and Living Image 4.3.1 software (PerkinElmer) set (excitation/emission: 420 nm/520 nm). The optical luminescent image data was displayed in pseudocolor that represents intensity in terms of Radiance (photons/sec/cm^2^/steradian) and calculated as average radiance per leaf.

### 4.6 *In vitro* transcription and infection

2μl of ToMV pTLW3 plasmid, ToMV^MP-ToBRFV^ or ToMV^MP-ToBRFV(1-216)^ were linearized with SmaI and cleaned by Gel extraction kit (Zymo Research, ZR-D4002). *In vitro* transcription using the mMESSAGE mMACHINE™ T7 kit (Invitrogen by Thermo Fisher Scientific, AM1344) was performed according to manufacturer’s instructions.

Two drops of 2.5μl of transcript were used to mechanically inoculate *N. benthamiana* plants that were dusted with carborundum powder prior to inoculation. Leaves of the infected plants were collected after 5-7 dpi to serve as inoculum for tomato plants.

### 4.7 Evaluation of ToMV levels by RT-qPCR Analysis

Total RNA was extracted from 50-100 mg of leaf tissue by Plant Total RNA Mini Kit (Geneaid, RPD050), which includes a DNAse treatement. RNA (500 ng) served as a template for cDNA synthesis (PCRBIO, PB30.11). RT-qPCR was performed using 2xSyGreen Mix (PCRBIO, PB20.16) in a Rotor-Gene 6000 cycler (Corbett Research). ToMV levels were calculated as the expression ratio of ToMV-CP and the gene of reference, TIP41 (Lacerda *et al.*, 2015).

### 4.8 Evaluation of ToMV and ToBRFV levels by SDS-PAGE and Western blot

For detection of the spread of the virus in tomato, the first small leaf (1-3 cm) and apex were collected and ground while frozen in a microcentrifuge tube. Laemmli buffer (40 μl of 3x) (100 mM Tris, 2% SDS, 20% glycerol, 4% β-mercaptoethanol, pH 6.8) were added and mixed into the sample, followed by 10 min centrifugation and 5 min boiling of the supernatant. Samples were run on 12% SDS-PAGE acrylamide gels, transferred to nitrocellulose membranes (Protran, #10401380) and blocked with 3% skimmed milk in Tris buffer saline (TBS)-Tween. CP of the virus was detected by rabbit anti-Tobamovirus CP; 1:20,000 (courtesy of Dr. Aviv Dombrovsky), and anti-Rabbit HRP; 1:20,000 (Jackson Immunoresearch, #323-005-021.). Chemiluminescence was observed using Elistar Supernova as substrate (Cyanagen, #XLSE2) and images of protein bands were acquired and quantified using the Alliance UVITEC software.

### 4.9 Cell-to-cell movement analysis

Constructs expressing MP-YFP and ER-mCherry were agroinfiltrated into *N. benthamiana* and visualized 36 hours later. For confocal imaging, we used an Olympus IX 81 inverted laser scanning confocal microscope (Fluoview 500) equipped with an OBIS 488 and 561 nm laser lines and a 60× 1.0 NA PlanApo water immersion objective. YFP and mCherry were excited at 488 and 561 nm and imaged using BA505-525 nm and BA575–620 nm emission filters respectively. The number of YFP-fluorescent cells surrounding the original cell of expression was quantified to determine the level of movement for each MP.

## Acknowledgements

We would like to thank Prof. Manfred Heinlein (CNRS, France), Prof. Bryce Falk (UC Davis, USA) for providing the TMV-GFP plasmid and Prof. Masayuki Ishikawa (NARO, Japan) for kindly providing the ToMV infectious clone. We also thank Dr. Aviv Dombrovsky for anti-tobamovirus CP antibody and Prof. Ilan Levin (ARO, Israel) for the tomato (cv. Moneymaker) lines used in the manuscript. We thank Dr. Amit Gal-On (ARO, Israel), Dr. Victor Gaba (ARO, Israel) and Dr. Munir Mawassi (ARO, Israel) for their valuable advice and critical reading of this manuscript. This work was funded by grant number 20-02-0130 from the Chief Scientist, Israeli Ministry of Agriculture.

## Conflict of interests

The authors declare no conflict of interests.

## Author contributions

HH and ZS designed the research, conducted the experiments and wrote the manuscript.

## Data availability statement

The data that support the findings of this study are available from the corresponding author upon reasonable request.

## Notes

### Competing Interest Statement

The authors have declared no competing interest.

